# Cooperative and aggressive behaviours vary between ranks in anemonefish social hierarchies

**DOI:** 10.1101/2021.01.19.427348

**Authors:** T Rueger, SJ Heatwole, MY Wong

## Abstract

Many animal groups consist of individuals organised in dominance hierarchies, based on age, size or fighting ability. Lower ranked individuals often do not reproduce themselves but perform cooperative behaviours to help the reproductive output of dominant individuals or the group as a whole. Theoretical models suggest that individuals of higher rank should show increased amounts of aggressive behaviours, such as aggressions towards other group members, but should decrease the amount of cooperative behaviours, such as brood care or territory maintenance. Most empirical tests of these models focus on insect or mammalian systems where kin selection plays a large role, rather than animals that live in groups of unrelated individuals. Here we use two anemonefish species to test hypotheses of variation in cooperation and aggression with respect to social rank and species, for social systems where group members are unrelated. We assessed the behaviours of each rank in 20 groups of *Amphiprion percula* and 12 groups of *A. perideraion* in Kimbe Bay, Papua New Guinea. We also performed a removal experiment to test if cooperative and aggressive behaviours are likely adaptive, i.e., if they change as an individual ascends in rank. Our results show differences between the two species, with *A. percula* showing more cooperative behaviours and *A. perideraion* showing more aggressive behaviours, despite them being closely related and sharing a very similar ecology. With respect to both cooperation and aggression we found consistent differences between ranks in both species, with higher ranks performing more aggressive as well as more cooperative behaviours. When we experimentally provided lower ranked individuals (rank 4) an opportunity to ascend in the hierarchy, they showed more aggression and more cooperation in line with our observations for rank 3 individuals. Thus, we show that rank specific behavioural patterns are likely adaptive in anemonefishes and that some model predictions do not hold in systems where kin selection benefits are absent. Rather, future fitness benefits through territory inheritance and group augmentation likely motivate cooperative and aggressive behaviours by subordinates in groups of unrelated vertebrates.

## Introduction

In many animal societies, groups are organized into dominance hierarchies or queuing systems, based for example on size, fighting ability or age (Cant, 2000; Shreeves & Field, 2002; Buston, 2003a). Although lower ranked individuals are often not part of the current breeding effort, they are generally capable of reproduction in some taxa, which begs the question of why they tolerate breeding suppression and cooperate. One explanation for this is that subordinates may gain indirect fitness benefits by helping to raise related offspring (Hamilton, 1963; 1964; West-Eberhard, 1975; Queller, 1994). Another reason why lower ranked individuals do not breed is that they have few chances to leave their natal territories and go elsewhere due to harsh environmental conditions or lack of breeding territories (Emlen, 1982), and while subordinates often have the capability to breed themselves, they may not attempt to breed within their group and instead opt to cooperate because of social constraints, such as the threat of punishment (Koenig & Pitelka, 1979; Cant et al., 2010). Additionally, subordinates may eventually inherit the breeding territory and thus gain future fitness benefits by improving the quality of the territory or promoting group augmentation (Wolfenden & Fitzpatrick, 1978; Kokko & Johnstone, 1999; Kokko et al., 2001). All of these aspects contribute to the evolution of cooperation and complex societies (reviewed in Bourke, 2011). However, what we do not fully understand is why cooperative effort often varies quite dramatically between individuals, even between those within the same group in the same species (Balshine et al., 2001; Stiver et al., 2006; Clutton-Brock et al., 2008; Wong, 2011).

Theory suggests that individuals in dominance hierarchies should reduce the amount of cooperation and increase the amount of aggression they perform when they are of higher rank and in larger groups (Cant & Field, 2001; 2005; Cant et al., 2006a; Field et al., 2006). According to several theoretical models, higher ranked subordinates in social groups have a higher probability of future fitness returns from inheriting the dominant position and should therefore put less effort into cooperative behaviours helping the dominants (Cant & Field, 2001; 2005; Field et al., 2006), and more effort into aggressive behaviours defending their rank (Cant et al., 2006a). However, these models primarily consider kin selection benefits from helping related group members and the associated future fitness benefits from inheriting dominant status in such groups. While kin selection predicts that the degree of relatedness will modify helping effort, several examples exist where relatedness does not explain variability in cooperative behaviour, e.g., in cichlid fishes (Le Vin et al., 2011) and some bird species (Wright et al., 1999; Canestrati et al., 2005). Models have thus far rarely considered future fitness benefits from group augmentation effects of cooperation, where individuals survive or reproduce better in larger groups (Kokko et al. 2001). These effects may be especially important in explaining cooperative behaviours in animals that live in groups of unrelated individuals (Kokko et al., 2001; Kingma et al., 2014). Also, Cant & Field (2005) show that model predictions do not hold when the cost of helping is higher in lower ranks, which may be common in vertebrates. Besides providing a conceptual underpinning for variation in cooperative behaviours among ranks, empirical tests of theoretical predictions have so far been conducted mainly in insect and mammalian groups where relatedness plays a large role (e.g., Cant et al., 2006b; Cronin & Field, 2007; Amsalem & Hefetz, 2011; Thavaraja et al., 2014, Jandt et al., 2014).

In vertebrates with dominance hierarchies where relatedness does not play a large role, such as many fish species, variation in cooperative or aggressive behaviours more likely depend on dominance rank and group size. Aggressive behaviours in some fish groups have been found to vary with dominance rank (Fricke & Fricke 1977, Colleter & Brown, 2011; Dey et al. 2012), although rank specific studies are still rare. In male rainbowfish (*Melanotaenia duboulayi*) various behaviours have been found to covary with the individual’s position in the size-based dominance hierarchy, with dominant fish being more aggressive, bolder and more active (Colleter & Brown, 2011). Cooperative behaviours by subordinates in complex fish groups have primarily been studied in cooperative breeding freshwater cichlids (Taborsky & Limberger, 1981), where different sized subordinates have been found to be task specialized, e.g., smaller subordinates performed egg-predator defence more frequently, whereas larger subordinates spend more time digging sand from the breeding shelter (Bruintjes & Taborsky, 2011). While most group-living marine fishes do not provide alloparental brood care, other behaviours beneficial to the group or the territory have been observed in several taxa, such as defence against egg predators, territory maintenance and cleaning (Ross, 1978; Fricke, 1979; Iwata & Manbo, 2013), though these behaviours have rarely been quantified or studied for specific ranks. Thus, quantifying cooperative behaviours as well as aggression in a wider range of vertebrate species, and considering the two behaviour types at the same time will give a better understanding about the differences between ranks in dominance hierarchies and thus broaden our understanding of sociality in general.

Here we use anemonefishes (genus Amphiprion) to examine differences in cooperative and aggressive behaviours among ranks and to experimentally test whether these differences are likely to be adaptive, i.e., whether there are predictable changes in behavioural patterns as an individual changes rank. Anemonefishes form symbiotic relationships with their host anemone (Fautin, 1992). They are sequential hermaphrodites and form strict size hierarchies within groups of unrelated individuals, where groups typically consist of one dominant breeding pair and zero to five unrelated subordinates (Fricke, 1979; Fautin, 1992; Buston, 2004a; Buston & Cant 2006). The female is always the largest individual (rank 1), the male is the second largest (rank 2), and all other fish are non-breeders that get progressively smaller (rank 3, rank 4, etc.) (Fricke & Fricke, 1977; Buston, 2003a). Rank ascension only happens when a higher rank disappears (Buston, 2003a; Buston, 2004b). Although subordinate anemonefishes are not known to help dominants rear offspring, alternate metrics of cooperation, namely territory defence (chasing predators away from the group) (Ross, 1978; Fricke; 1979; Iwata & Manbo, 2013) and territory maintenance (cleaning the anemone) (Mariscal 1966) can be observed and quantified. Specifically, we focused on two closely related anemonefish species common in the Indo-Pacific; *Amphiprion percula* and *A. perideraion*. While the two species both primarily inhabit the same anemone species, *Heteractis magnifica*, and are otherwise ecologically similar, differences in their behaviours have recently been reported. *A. perideraion*, particularly individuals at the top of the dominance hierarchy, have been observed leaving their anemone and forcefully taking over groups of *A. percula* (Rueger et al., 2018). In contrast, *A. percula* has been shown to not leave the confines of their host, even to bridge small distances (Branconi et al., *in press*). Thus, there is the potential for different costs and benefits of within-group rank ascension, making these two species appropriate for testing the robustness of current theory.

## Hypotheses

To investigate whether and how cooperative and aggressive behaviours vary between social ranks, we used a combination of field observations and experimental techniques to test the following specific hypotheses in *A. percula* and *A. perideraion*: 1) Cooperative and aggressive behaviours will differ depending on individual dominance rank but not species; 2) In line with current theoretical predictions, the frequency of cooperative behaviours will decrease and the frequency of aggressive behaviours will increase when individuals are experimentally promoted in rank.

## Methods

The study was conducted on inshore reefs near Mahonia Na Dari Research and Conservation Centre in Kimbe Bay, Papua New Guinea (5°30’ S, 150°05’ E), in September and October 2018 and May 2019. All fieldwork was conducted using SCUBA.

### Study organisms

We used 20 groups of *A. percula*, and 12 groups of *A. perideraion* for observations and experiments. The groups were found on seven different inshore reefs. Out of the 20 *A. percula* groups, most (n=16) were associated with the anemone species *Heteractis magnifica*, while the rest were associated with *Stichodactyla gigantea* (n=4). This accurately reflects the observed relative distribution of host occupancy for the population on inshore reefs in Kimbe Bay (Chausson et al. 2018). All *A. perideraion* groups (n=12) were associated with *H. magnifica* hosts. For *A. percula,* groups comprised four fish (n=16) or five fish (n=4). For *A. perideraion,* groups comprised either four fish (n=9) or five fish (n=3). The difference in sample size between species was due to a lack of available groups of *A. perideraion* with 4-5 individuals. Half of the *A. percula* groups (n=10) were observed breeding during the experiment, whereas few groups of *A. perideraion* were observed breeding (n=3).

### Rank ascension experiment

To experimentally examine variability in cooperative and aggressive behaviours between ranks we removed the rank 3 individual from half of the groups of *A. percula* (n=10) and *A. perideraion* (n=6) (i.e., treatment groups; Figure 1) to give the rank 4 an opportunity for rank ascension. From the other half of the groups (i.e., control groups, Figure 1) we removed the lowest ranking individual (rank 4 or 5 depending on the group size). This was done to control for the reduced number of group members and to be able to compare rank 4 behaviours from the treatment to rank 3 behaviours from the control groups. During the experiment, the removed individuals were held in aerated sea water tanks at Mahonia Na Dari Research and Conservation Centre, where they were provided with shelter and fed twice a day with dried brine shrimp (Omega One, freeze dried brine shrimp) and fish pellets (New Life Spectrum, marine fish food 1 mm pellets). The removed fish was then replaced back into its original group. Groups were filmed with a GoPro Hero5 for 12 minutes from a 1.5-2.5m distance at six time points: 1. Prior to removal of the rank 3 (or rank 4 in control groups), 2. Immediately after removal of the fish, 3. 24 hrs post-removal, 4. 48 hrs post-removal, 5. Immediately after the reintroduction of the removed individual, 6. 24 hrs post-reintroduction. To account for any ecological effects of anemone size, soft tailor’s tape was used to measure the long and short diameter of the anemone’s tentacle crown. Anemone size was then calculated as the area of an ellipse (π × major axis/2 × minor axis/2).

**Figure 1.**
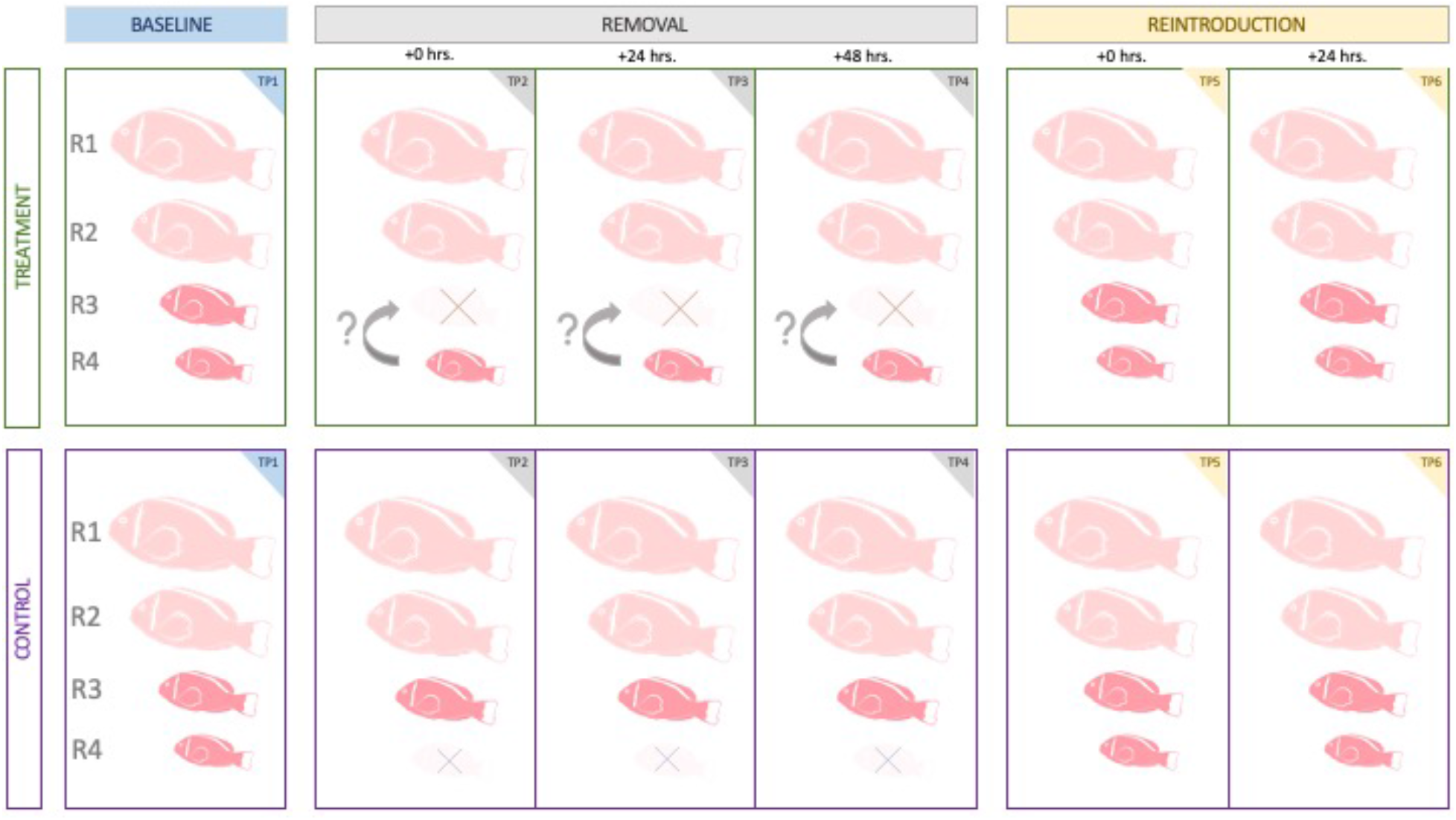
Experimental design of the rank ascension experiment conducted with groups of *A. percula* (n=20) and *A. perideraion* (n=12) in Kimbe Bay, Papua New Guinea. In treatment groups, the rank 3 (R3) was removed allowing for rank ascension by rank 4 (R4). In control groups, the lowest ranked group member was removed to control for group size reduction whilst preventing rank ascension by any group member. TP= Time point. R= rank.

### Video analysis

A total of 153.8hrs of video footage was analysed. Behaviours were scored using the BORIS program (Friad & Gamba, 2016) and following an ethogram based on Wong et al. (2013) (Supplemental Material Table 1). For each video, 10 of the 12 minutes of footage was watched with 1 minute at the start and end of each video being discarded to account for disturbance created by the diver approaching to handle the video camera. To account for differences in the time each fish was visible in the frame, the time each fish spent out of sight during the recording was noted and all behaviours were analysed per minute of observation. Each video was watched multiple times so that every individual in the group was scored separately and continuously in one sequence to make behaviour scores as accurate as possible. Behaviours scored fell into four broad categories: 1) aggressive, 2) submissive, 3) neutral, 4) cooperative (Table 1). For all interactive behaviours, the individual receiving the submission or aggression was recorded. Behaviours were scored for ranks 1 to 5.

**Table 1.** Broader behavioural categories used in the analysis for *A. percula* and *A. perideraion*. For a more detailed description of behaviours refer to the ethogram in supplemental material.

| Behavioural category | Description |
| --- | --- |
| Aggression | All aggressive behaviours among group members, e.g., chases, bites. |
| Submission | All submissive behaviours among group members, e.g., body shake. |
| Neutral | When group members were within one body length of each other but did not display aggression or submission. |
| Cooperation | Behaviours beneficial to the group (e.g., aggression towards egg predators, aggression towards food competitors, parental care) or beneficial to the anemone (e.g., anemone cleaning/manipulating, aggression towards anemone predators) |

### Data analysis

We used R (R Core Team, 2019) and the *lme4* package (Bates, Maechler & Bolker, 2012) to perform linear mixed model analysis (LMM). We used Akaike Information Criterion (AIC) (Sakamoto et al., 1986) for model selection. If ∆AIC was below 2, a likelihood ratio test was performed to decide which model was the best fit, with preference for the simpler model. Significance tests for LMMs were performed by likelihood ratio tests of the full model with the effect in question against the model without the effect. No obvious deviations from normality and homoscedasticity were detected by visually inspecting the residual plots. Models were also tested for outliers and collinearity between variables with *performance* (Luedecke et al., 2020). No outliers or high variance inflation factors (VIF) were detected in any of the best-fit models. Conditional and marginal R^2^ were calculated using Nakagawa’s R^2^ in *performance* (Nakagawa & Schielzeth, 2013; Johnson, 2014; Nakagawa, Johnson & Schielzeth, 2017).

As response variables, behaviours were analysed as behaviours per minute of observation to account for differences in observation time for each fish. Behaviours/min was log transformed to account for a non-normal distribution (left screw due to many zero values).

#### Behavioural differences between ranks, species and group sizes

To test the differences in behaviours between ranks and species, we fitted four separate LMMs for each behavioural category: aggression, submission, neutral behaviours and cooperation (Table 1). The corresponding response variables were aggression/min, submission/min, neutral/min and cooperation/min. For cooperation/min we performed model selection excluding parental behaviours (see Suppl. Table 1 for definition), since those were only recorded in *A. percula*. Fixed factors tested were fish species (*A. percula, A. perideraion*) and individual rank in the group hierarchy (ranks 1-5). Covariates tested included number of fish in each group (group size), anemone species (*H. magnifica, S. gigantea*) and anemone size (tentacle crown surface area of the anemone in cm^2^). Model selection was performed as described above. Group ID was used as a random variable to account for non-independence of individual behaviours within the same group. In the model with submission/min as the response variable, the random term variance component was zero and conditional R^2^ could not be calculated. This had no effect on the model components reported here (Pasch, Bolker, & Phelps, 2013). To be consistent with the other models in this section and to not influence AIC values for the model selection process, we left the random variable in the model.

#### Rank ascension experiment

To test differences in behaviours in the treatment and control groups and between different time points (Figure 1), separate LMMs were fitted for each species (*A. percula, A. perideraion*) and each rank in the group hierarchy. Since rank 3 individuals were absent from time points 2-4 in the treatment groups and rank 4 individuals were absent from time points 2-4 in the control groups, the full cross model (including rank, treatment group and time point as fixed factors and interactions) was only used to examine specific contrasts (see below). For the species and rank specific models, only ranks 1-4 were tested since there was limited data available for rank 5. The three behavioural categories of primary interest were used as response variables (aggression/min, submission/min, cooperation/min). Fixed factors tested were time point (time points 1-6 of the experiment, Figure 1), and treatment (treatment, control). Covariates tested were group size, anemone size and anemone species. Model selection was performed as described above. Group ID was used as a random variable to account for repeated measures of the same individuals. In two models (*A. perideraion*, rank 3, aggression/min and submission/min), the random term variance component was zero and conditional R^2^ could not be calculated. This had no effect on the model components reported here (Pasch, Bolker, & Phelps, 2013). To be consistent with the other models in this section and to not influence AIC values for the model selection process, we left the random variable in the model.

To further investigate variability due to rank, we compared the behaviour of ranks 3 and 4 directly in two ways. First, we compared the behaviour of the rank 3 in control groups (rank 4 removed) to the behaviour of the ascended rank 4 in treatment groups (rank 3 removed) using post-hoc tests on the difference in behaviours between time points using the *emmeans* package (Length, 2020). Second, we compared rank 3 behaviour during the baseline time point 1 (in control and treatment) and during the time points 2-4 (in control) to rank 4 behaviour during time points 2-4 (treatment). This comparison is important because during the removal, rank 3 in the control groups and rank 4 in the treatment groups effectively occupy the same position in the hierarchy (position 3). For this comparison, we built a full cross model (including all two-way and three-way interactions) with rank, treatment and time point as fixed factors, and group ID as a random effect. The model was checked for fit, collinearity and outliers as described above. We then used *emmeans* to build a custom contrast matrix for the above-mentioned comparisons. P-values were adjusted for multiple testing using a multivariate *t* distribution correction.

## Results

### Behavioural differences between ranks and species

#### Aggression

There were significant differences in the frequency of aggressive behaviours between ranks (χ_(4)_ = 145.080, p < 0.001), species (χ_(1)_ = 9.540, p = 0.002), and group sizes (χ_(1)_ = 14.730, p < 0.001) (Figure 2). The fixed effects explained 22.4% of the variance (R^2^_c_ = 0.254, R^2^_m_ = 0.224). The best fit model did not include anemone size or anemone species, or interactions between any fixed effects. The model estimates show that for each species, ranks 1-3 were more aggressive than the lower ranks (Figure 3a), and on average, rank 4 showed 33% (± 5% s. e.) less aggression than rank 1. An increase in group size by one fish increased mean aggressions per minute by 9% (± 2% s. e.). Looking specifically at how aggressive behaviours were directed towards each group member, ranks 1 and 2 in *A. perideraion* groups were more aggressive towards lower ranks than were ranks 1 and 2 in *A. percula* groups (Figure 4). Overall, aggression was 14% (± 5% s. e.) higher in *A. perideraion* compared to *A. percula* (Figure 2).

**Figure 2.**
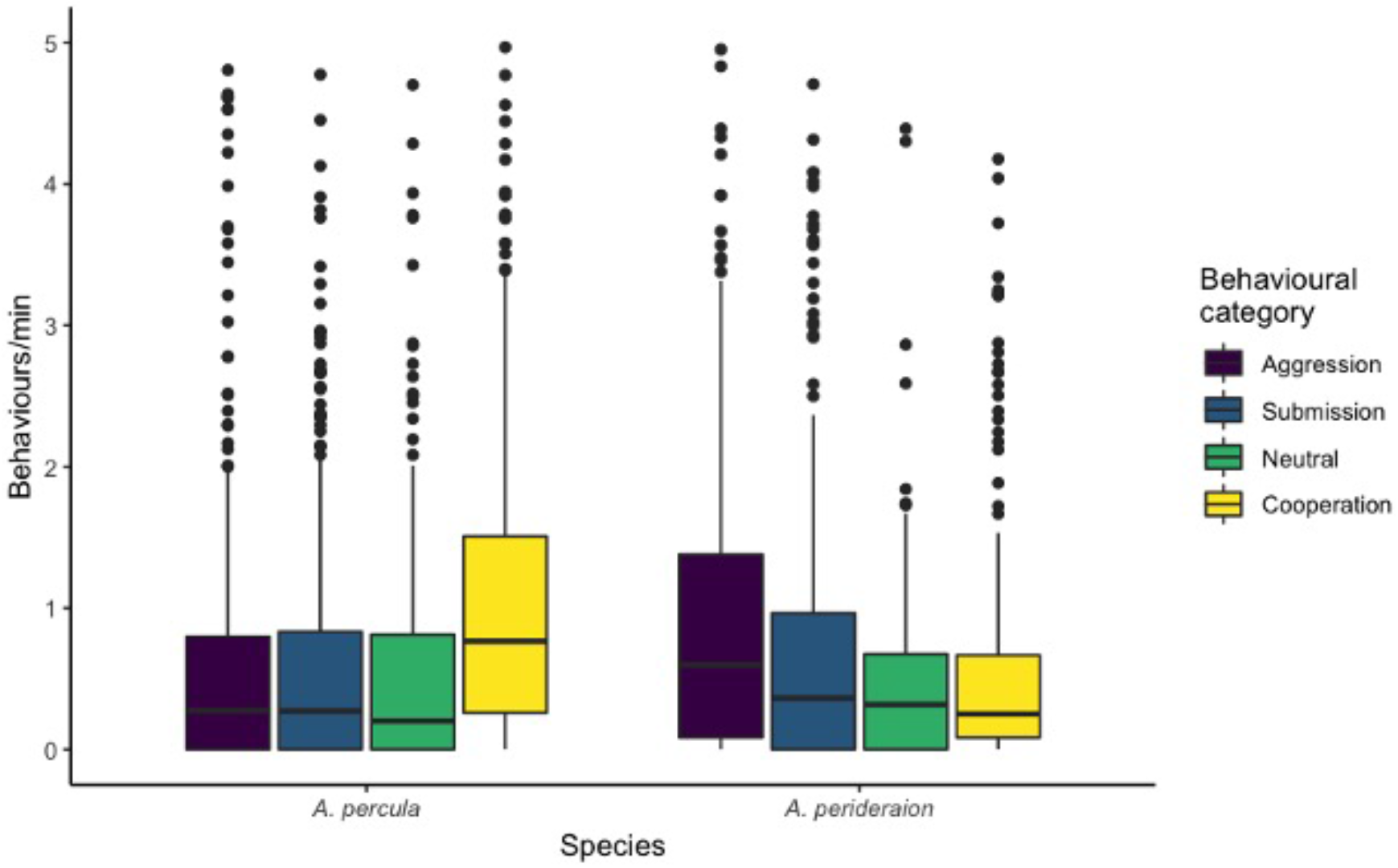
Behavioural frequencies (behaviours/min) for two species of anemonefishes (*Amphiprion percula* and *A. perideraion*) in Kimbe Bay, Papua New Guinea (central bar: median; boxes: lower and upper quartiles; whiskers: +/−1.5*IQR (interquartile range)). Observed behaviours were pooled into four categories (see Table 1).

**Figure 3.**
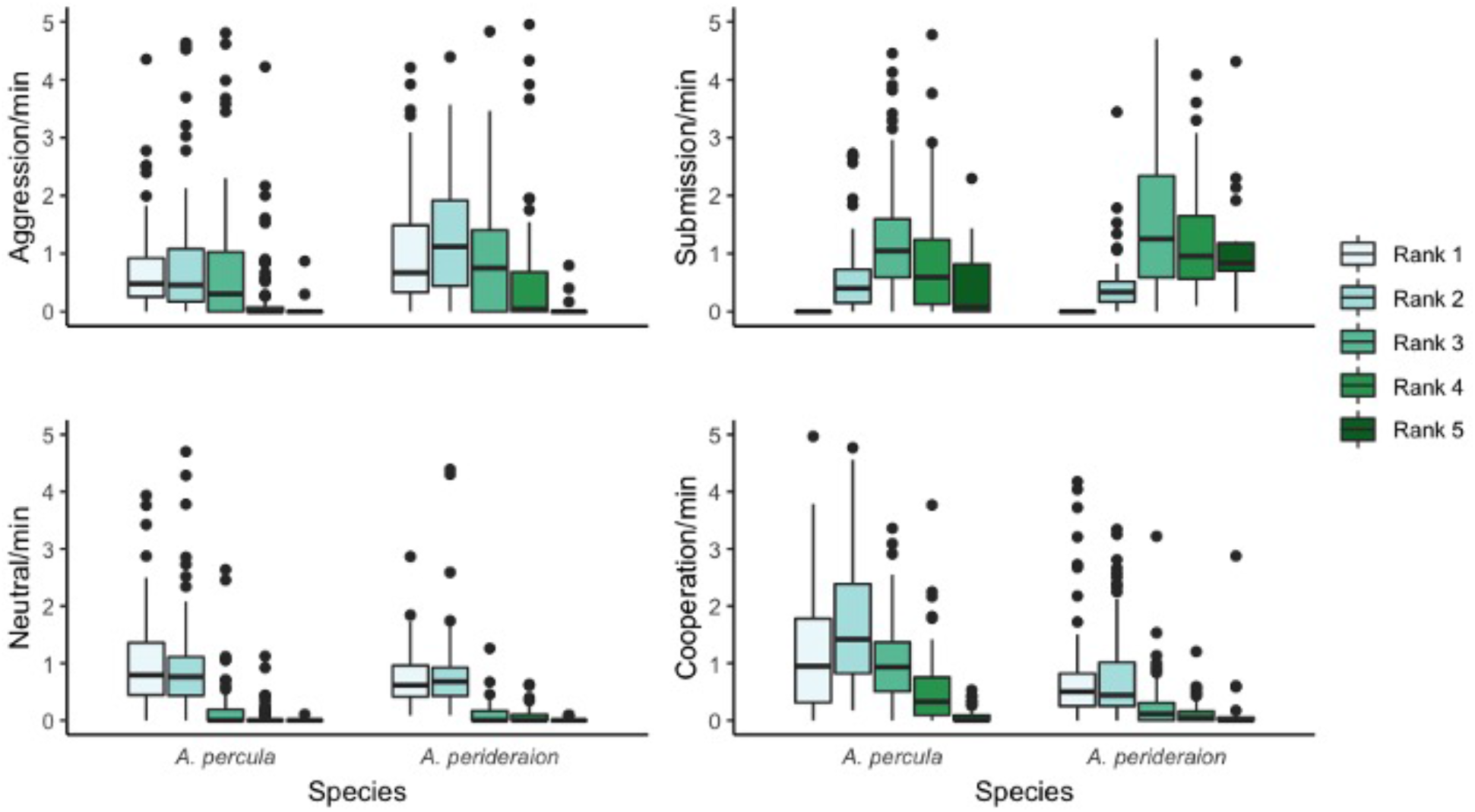
Behavioural frequencies (behaviours/min) for each rank (1-5) in the size hierarchy in two species of anemonefishes (*Amphiprion percula* and *A. perideraion*) (central bar: median; boxes: lower and upper quartiles; whiskers: +/−1.5*IQR (interquartile range)). Observed behaviours were pooled into four categories (see Table 1); a) Aggression; b) Submission; c) Neutral; d) Cooperation.

**Figure 4.**
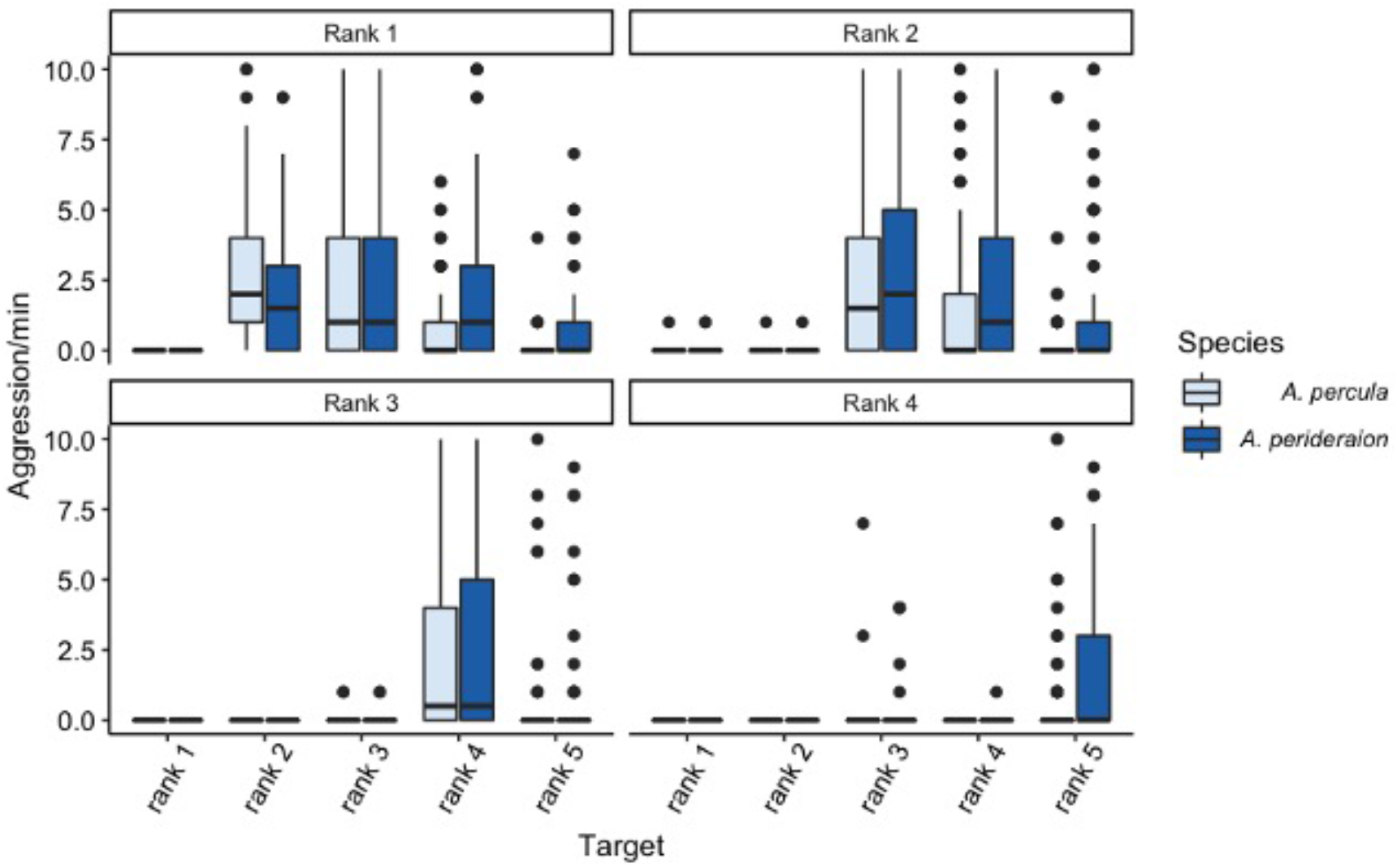
Number of aggressive behaviours per minute (Aggression/min) recorded for rank 1 to rank 4 in two species of anemonefishes (*Amphiprion percula* (n=20) and *A. perideraion* (n=12) (central bar: median; boxes: lower and upper quartiles; whiskers: +/−1.5*IQR (interquartile range)). The individual the aggression was directed towards was recorded (ranks 1-5).

#### Submission

There were significant differences in the frequency of submissive behaviours between ranks (χ_(4)_ = 331.44, p < 0.001). The fixed effects explained 34.7% of the variance (R^2^_c_= NA, R^2^_m_= 0.347). The best fit model did not include anemonefish species, group size, anemone size or anemone species, or any interactions. The most submissive behaviours were performed by rank 3 and the least by rank 1, with a mean difference in submission frequency between the two of 86% (± 5% s.e.) (Figure 3b). Most submissions by ranks 2-4 were performed towards the rank immediately above them (*A. percula*: rank 2 to rank 1, 100%, rank 3 to rank 2, 53%, rank 4 to rank 3, 58%; *A. perideraion*: rank 2 to rank 1, 100%, rank 3 to rank 2, 53%, rank 4 to rank 3, 63%).

#### Neutral

There were significant differences in the frequency of neutral social behaviours between ranks (χ_(4)_ = 497.57, p < 0.001). The fixed effects explained 45.9% of the variance (R^2^_c_= 0.505, R^2^_m_= 0.459). The best fit model did not include anemonefish species, group size, anemone size or anemone species, or any interactions. Neutral behaviours among group members were primarily observed between rank 1 and rank 2 in both species and consisted primarily of close proximity (Figure 3c).

#### Cooperation

There were significant differences in the frequency of cooperative behaviours between ranks (χ_(4)_ = 140.950, p < 0.001) and species (χ_(1)_ = 12.753, p < 0.001). The best fit model did not include group size, anemone size, anemone species or any interactions. The fixed effects explained 20.5% of the variance (R^2^_c_ = 0.263, R^2^_m_ = 0.205). Cooperative behaviours were observed at a 26% (± 8% s.e.) higher rate in *A. percula* compared to *A. perideraion* (Figure 2). For each species, ranks 1-3 were more cooperative than the lower ranks (Figure 3d). The specific cooperative behaviours performed also differed between ranks (Figure 5). For example in *A. percula*, interaction with the anemone (‘cleaning’) made up approximately 39% of cooperative behaviours for ranks 1 and 2, whereas for ranks 4 and 5 this made up the majority of cooperative behaviours (53% and 75% respectively) (Figure 5).

**Figure 5.**
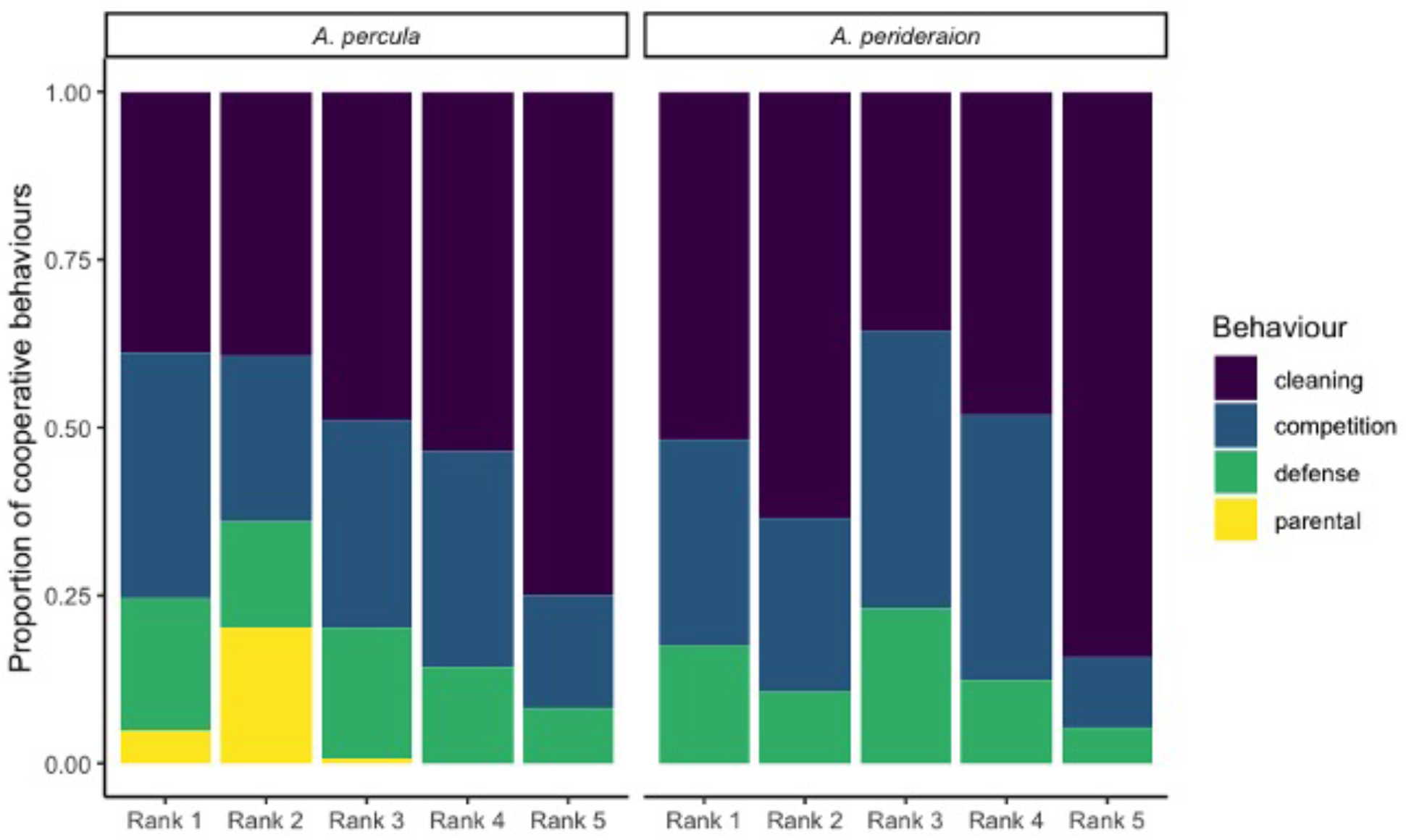
Proportion of cooperative behaviours for each rank in *A. percula* and *A. perideraion*. Behaviours include ‘cleaning’ (interactions with the anemone including cleaning, massaging, biting), ‘competition’ (aggressive behaviours towards food competitors such as other Pomacentridae), ‘defense’ (aggressive behaviours towards egg predators such as Labridae or anemone predators such as Chaetodontidae), and ‘parental’ (when the fish is within one body length of an egg clutch and is fanning or mouthing the brood).

### Rank ascension experiment

#### Aggression

Out of 32 removed fish, none were evicted from their groups upon reintroduction. All reintroduced fish received aggression initially, typically by all group members (see below), but were allowed to remain close to the anemone.

For both species, the frequency of aggressive behaviours towards other group members differed between time points of the experiment for all ranks tested (Table 2, Figure 6). For *A. percula*, the best fit model for ‘aggressions/min’ for ranks 1-3 only included time point as a fixed term, and did not include treatment, group size or any interactions. For *A. perideraion*, the best fit model for rank 1 and rank 2 only included time point as a fixed term, and did not include treatment, group size or any interactions. For *A. perideraion* rank 3, the best fit model also included group size, with aggressions rising by an estimated 7% (± 3% s. e.) for an additional group member. For rank 4 of both species, the best fit model included time point, treatment, group size, as well as an interaction between time point and treatment, all of which were significant (Table 2).

**Table 2.** Best fit models for each rank of *A. percula* and *A. perideraion* with aggressive behaviours towards other group members per minute (‘aggression/min’) as the response variable. Fixed terms tested included time point of the experiment (‘time point’), treatment group (‘treatment’), and interactions, with number of fish in each group (‘group size’) tested as a covariate. Group ID was included as a random variable to account for repeated measures. Significance was tested using likelihood ratio tests.

| *A. percula* |  |  |  | *A. perideraion* |  |  |  |
| --- | --- | --- | --- | --- | --- | --- | --- |
| Term<br>(log(aggression/min)<br>~) | DF | X<sup>2</sup> | p-value | Term<br>(log(aggression/min)<br>~) | DF | X<sup>2</sup> | p-value |
| **Rank 1** ($R^2_c = 0.347$, $R^2_m = 0.188$) | | | | **Rank 1** ($R^2_c = 0.383$, $R^2_m = 0.195$) | | | |
| time point | 5 | 30.795 | <0.001 | time point | 5 | 23.678 | <0.001 |
| **Rank 2** ($R^2_c = 0.193$, $R^2_m = 0.147$) | | | | **Rank 2** ($R^2_c = 0.336$, $R^2_m = 0.199$) | | | |
| time point | 5 | 20.599 | <0.001 | time point | 5 | 22.587 | <0.001 |
| **Rank 3** ($R^2_c = 0.567$, $R^2_m = 0.424$) | | | | **Rank 3** ($R^2_c = \text{NA}$, $R^2_m = 0.550$) | | | |
| time point | 5 | 58.581 | <0.001 | time point | 5 | 49.926 | <0.001 |
|  |  |  |  | group size | 1 | 5.554 | 0.018 |
| **Rank 4** ($R^2_c = 0.563$, $R^2_m = 0.530$) | | | | **Rank 4** ($R^2_c = 0.847$, $R^2_m = 0.840$) | | | |
| time point | 5 | 29.272 | <0.001 | time point | 5 | 41.794 | <0.001 |
| treatment | 1 | 7.889 | 0.005 | treatment | 1 | 24.738 | <0.001 |
| group size | 1 | 11.415 | 0.001 | group size | 1 | 28.227 | <0.001 |
| time point x treatment | 2 | 29.123 | <0.001 | time point x treatment | 2 | 63.685 | <0.001 |

**Figure 6.**
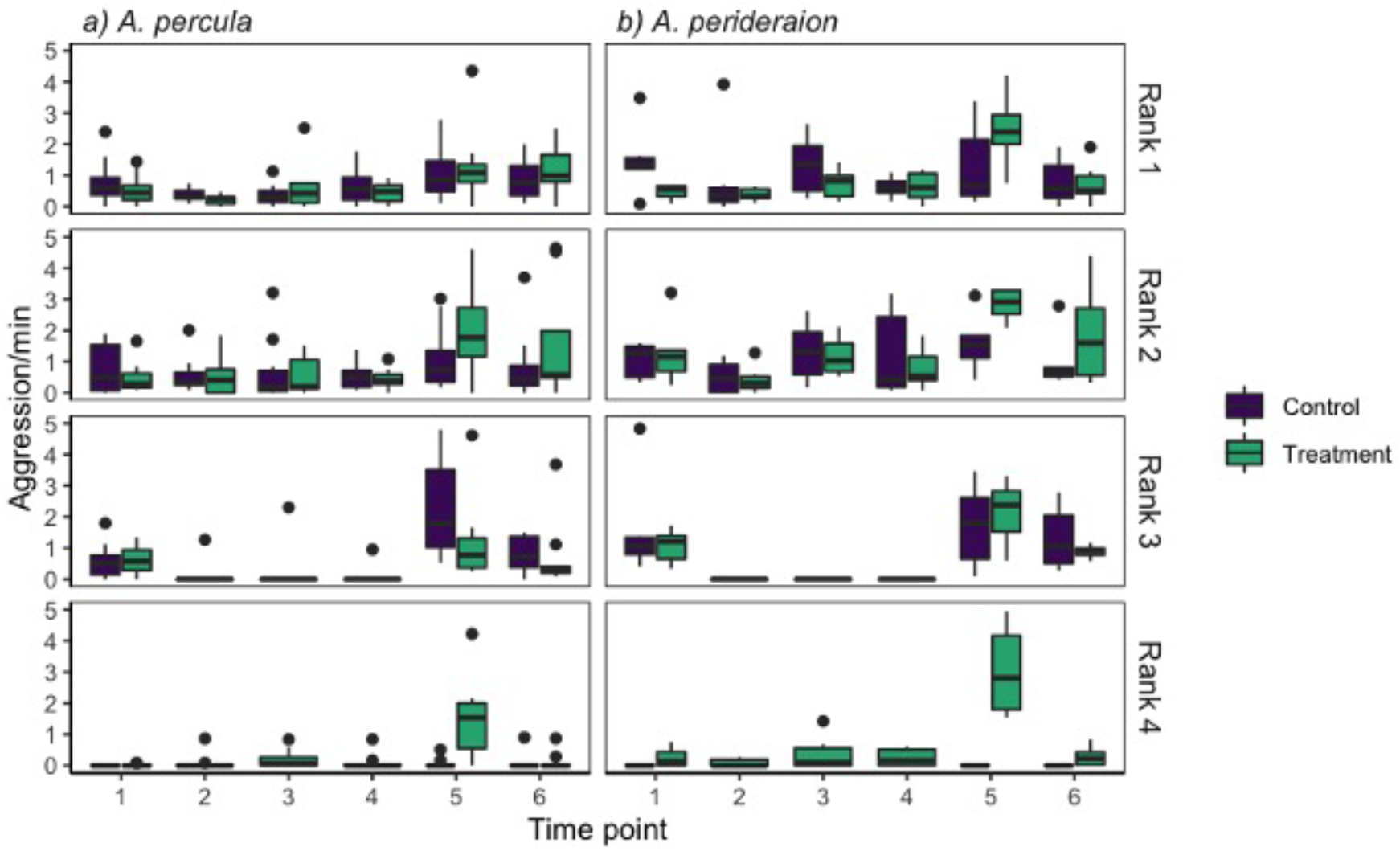
Aggressive behaviours per minute (‘Aggression/min’) in groups of a) *Amphiprion percula* (n=20) and b) *A. perideraion* (n=12), for each rank in the size hierarchy (rank 1-4), time point (1-6) and treatment group (purple: Control, green: Treatment) of the rank ascension experiment (central bar: median; boxes: lower and upper quartiles; whiskers: +/−1.5*IQR (interquartile range)).

In the control groups of both species, rank 3 showed a significantly reduced rate of aggression while rank 4 was removed (TP2-4) compared to the baseline time point (TP1) (Suppl. Figure 1, TP1 vs TP2, *A. percula*: estimate=-0.322, 95% confidence interval (c. i.)= −0.606, −0.038; *A. perideraion*: estimate= −0.809, c. i.= −1.341, −0.279). In the *A. percula* treatment groups, rank 4 showed slightly increased aggression while rank 3 was removed, though this was not significant, while in the *A. perideraion* treatment groups rank 4 showed no change (Suppl. Figure 1). Compared to the control groups rank 3 baseline time point (TP1 of the control), the rank 4 of the treatment groups showed a significantly lower frequency of aggression in both species (Suppl. Figure 1, rank 3 control TP1 vs rank 4 treatment TP2, *A. percula:* estimate=-0.385, c. i.= −0.718, − 0.051; *A. perideraion*: estimate=-0.717, c. i.=-1.101, −0.334). Levels of aggression in the treatment group rank 4 while rank 3 was removed were not significantly different compared to levels of aggression in the control group rank 3 while rank 4 was removed (Suppl. Figure 1).

The highest frequency of aggression towards other group members for all ranks in both species was recorded at time point 5, immediately after reintroduction (Figure 6). For this time point, the rank 4 in the treatment groups showed an estimated 67% (± 13% s. e.) higher rate of aggression in *A. percula*, and a 112% (± 14% s. e.) higher rate of aggression in *A. perideraion*, compared to the baseline time point 1 (Suppl. Figure 1). All group members directed a large proportion of aggressions towards the reintroduced fish at time point 5 in the treatment groups (immediately after rank 3 introduction) (*A. percula*: rank 1, 71%, rank 2, 84%, rank 4, 72%, *A. perideraion*: rank 1, 63%, rank 2, 83%, rank 4, 95%). In the control groups at time point 5, a large portion of aggressions were directed towards the newly reintroduced rank 4 compared to the baseline time point 1 (*A. percula*: rank 1: TP1, 17%, TP5, 24%; rank 2: TP1, 15%, TP5, 39%; rank 3: TP1, 100%, TP5, 97%; *A. perideraion*: rank 1: TP1, 42%, TP5, 60%; rank 2: TP1, 26%, TP5, 40%; rank 3: TP1, 84%, TP5, 96%).

#### Submission

For *A. percula*, the frequency of submissive behaviours by ranks 2 and 3 did not differ with treatment or time point of the experiment (Figure 7a). The best fit model for ranks 2 and 3 was the null model, with the random factor (Group ID) explaining 15% and 5% of the variance respectively (rank 2 R^2^_c_= 0.147, rank 3 R^2^_c_=0.049). For rank 4 *A. percula*, the best fit model included only time point as the fixed variable, which explained 23% of the variance (R^2^_c_= 0.259, R^2^_m_ = 0.232). For *A. perideraion*, the frequency of submissive behaviours did not differ with treatment or time point of the experiment for any rank (Figure 7b). The best fit model was the null model for ranks 2-4, with the random factor (Group ID) explaining 8% and 45% of the variance in rank 2 and 4 respectively (rank 2 R^2^_c_= 0.077, rank 3 R^2^_c_= NA, rank 4 R^2^c=0.449).

**Figure 7.**
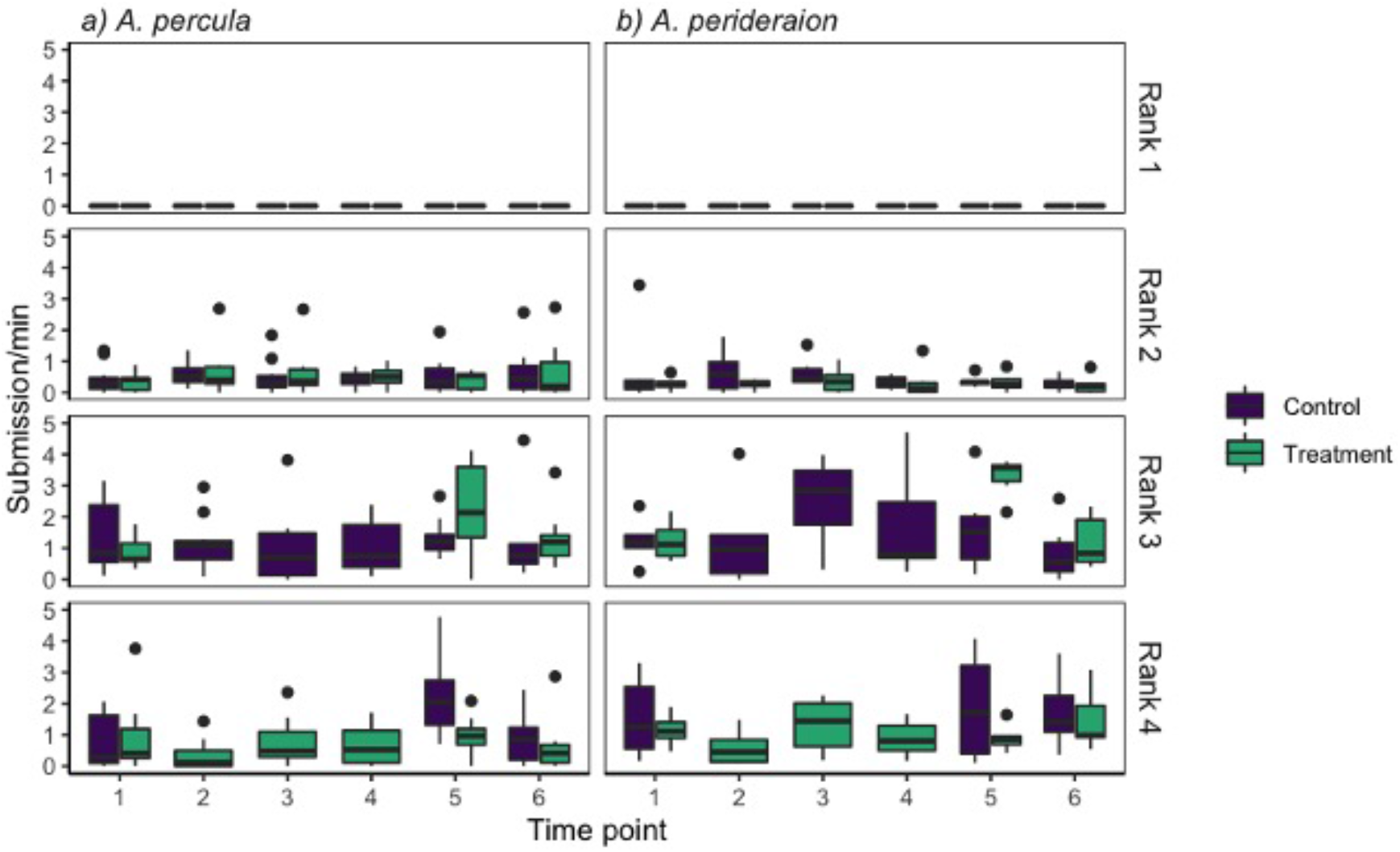
Submissive behaviours per minute (‘Submission/min’) in groups of a) *Amphiprion percula* (n=20) and b) *A. perideraion* (n=12), for each rank in the size hierarchy (rank 1-4), time point (1-6) and treatment group (purple: Control, green: Treatment) of the rank ascension experiment (central bar: median; boxes: lower and upper quartiles; whiskers: +/−1.5*IQR (interquartile range)).

For both species, there were no significant differences in submissive behaviours in rank 4 during the treatment (Suppl. Figure 2), but the main target of submissions changed from rank 3 at time point 1 (*A. percula* 53%, *A. perideraion* 66% of submissive behaviours) to rank 2 during the time rank 3 was removed at time points 2 (*A. percula* 67%, *A. perideraion* 63%), 3 (*A. percula* 56%, *A. perideraion* 58%) and 4 (*A. percula* 62%, *A. perideraion* 67%). In *A. percula*, at time points 5 and 6 in the treatment, rank 4 showed submissive behaviours towards ranks 1, 2 and 3 (TP5: 27%, 46% and 27% respectively; TP6: 30%, 30% and 40% respectively). In *A. perideraion*, at time point 5 in the treatment, rank 4 showed similar frequencies of submissive behaviours to rank 1 and 2 (44% and 39% respectively), and at time point 6, most were directed at rank 3 once again (70%). Rank 3 did not change the frequency or target of submissive behaviours during the control in either species (Suppl. Figure 2). Spikes in submissive behaviours were recorded for rank 4 and rank 3 when the fish were reintroduced at time point 5. At reintroduction, rank 4 *A. percula* performed a mean of 27% (± 8% s. e.) more submissions compared to time point 1 (Figure 7a). In *A. perideraion* overall, rank 4 showed less submissive behaviours in the treatment group than rank 3 in the control group during the early treatment time points, whereas later treatment time points were not significantly different from the rank 3 control (Suppl. Figure 2).

#### Cooperation

For ranks 1-3 in both species, the frequency of cooperative behaviours did not differ with treatment or time point of the experiment (Figure 8). The best fit model for ranks 1-3 was the null model, which in *A. percula* explained 29%, 38%, and 18% of the variance respectively (rank 1 R^2^_c_= 0.287, rank 2 R^2^_c_= 0.383, rank 3 R^2^c=0.184), and in *A. perideraion* explained 15%, 12%, and 3% of the variance respectively (rank 1 R^2^_c_= 0.148, rank 2 R^2^_c_= 0.124, rank 3 R^2^c=0.033). For rank 4 in both species, the best fit model included only treatment as the fixed variable. The frequency of cooperative behaviours per minute of observation for rank 4 was 21% (± 10% s. e.) higher in the treatment group compared to the control group in *A. percula* (χ_(1)_ = 4.394, p = 0.036), and 8% (± 4% s. e.) higher in the treatment group compared to the control group in *A. peridareion* (χ_(1)_ = 4.887, p = 0.027). ‘Treatment’ explained 5.6% of the variance in *A. percula* (R^2^_c_= 0.108, R^2^_m_ = 0.056) and 7.5% of the variance in *A. perideraion* (R^2^_c_= 0.080, R^2^_m_ = 0.075).

**Figure 8.**
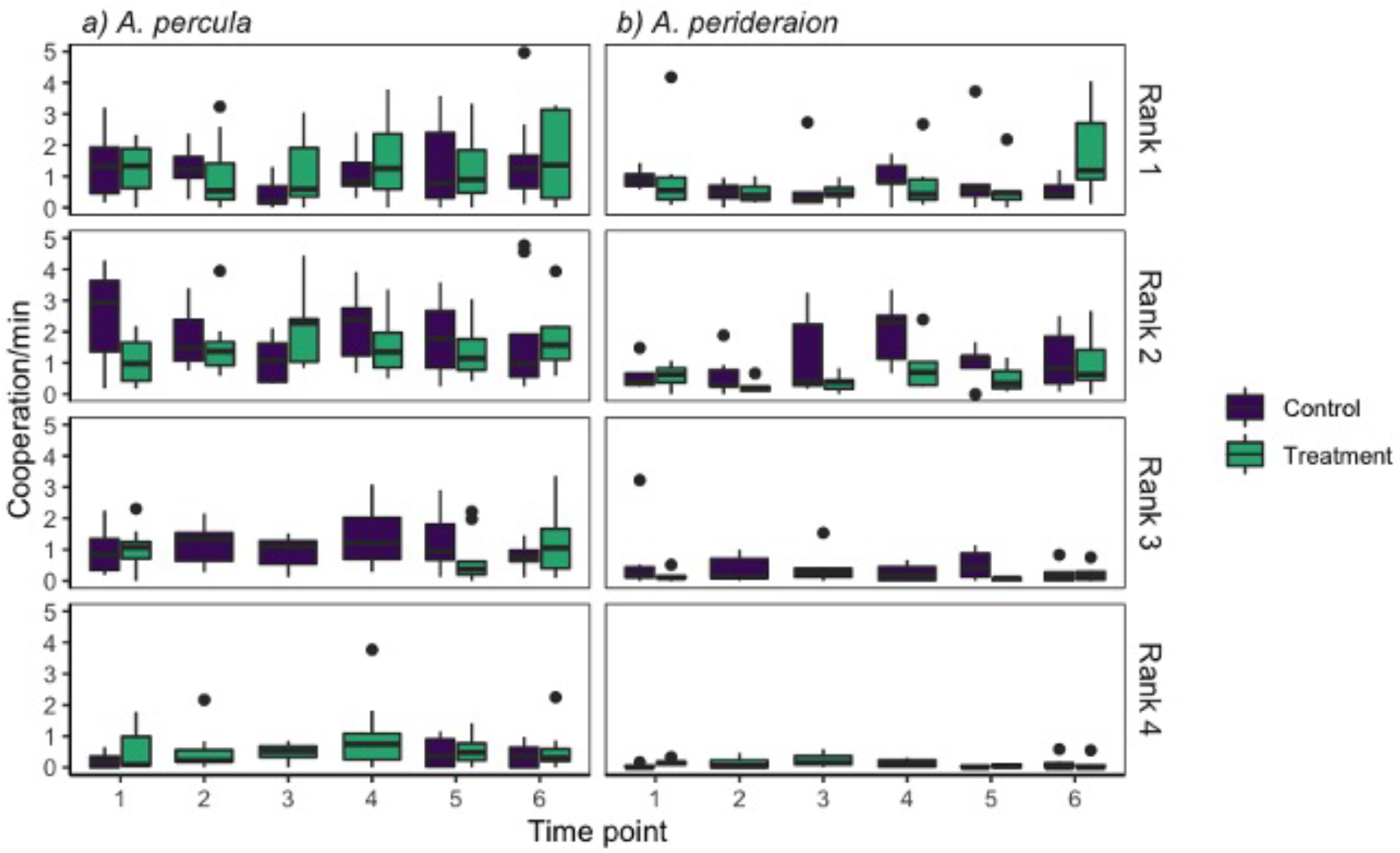
Cooperative behaviours per minute (‘Cooperation/min’) in groups of a) *Amphiprion percula* (n=20) and b) *A. perideraion* (n=12), for each rank in the size hierarchy (rank 1-4), time point (1-6) and treatment group (purple: Control, green: Treatment) of the rank ascension experiment (central bar: median; boxes: lower and upper quartiles; whiskers: +/−1.5*IQR (interquartile range)).

In *A. percula* cooperation in rank 4 was highest at time point 4 (48h after removal of rank 3), when cooperative behaviours were performed on average at a 2.8 times higher rate compared to the baseline time point 1 (mean cooperation/min ± s. e., time point 1: 0.35 ± 0.1; time point 4: 0.96 ± 0.3). In particular, ‘cleaning’ behaviours increased the most, which made up 30% of cooperative behaviours at baseline time point 1 and 50% at treatment time point 3. In *A. perideraion*, cooperation in rank 4 in the treatment was highest at time point 3 (24h after removal of rank 3), when cooperative behaviours were performed on average at a 1.8 times higher rate compared to the baseline time point 1 (mean cooperation/min ± s. e., time point 1: 0.21 ± 0.05; time point 3: 0.37 ± 0.16). The behaviour that increased the most for *A. perideraion* rank 4 was also ‘cleaning’, which made up 40% of cooperative behaviours at baseline time point 1 and 57% at treatment time point 3. In both species, post-hoc tests showed no significant differences between time points for the rank 4 in the treatment groups (baseline TP1 compared to TP2-4 when rank 3 is removed and rank ascension is possible) or the rank 3 in the control groups (baseline TP1 compared to TP2-4 when rank 4 is removed and group size is adjusted) (Suppl. Figure 3). In *A. percula*, cooperative behaviours were significantly lower in rank 4 during the early treatment time point 2, compared to the rank 3 baseline time point (TP1, estimate = −0.465, c. i.= −0.917, −0.012) and control time points (TP2, estimate=-0.47, c. i.=-0.922, −0.017, and TP4, estimate=-0.536, c. i.=-1.001, −0.072). In contrast, the frequency of cooperative behaviours of rank 4 during the later treatment time points (24h/48h after rank 3 removal, TP3/TP4), were not significantly different from the cooperations recorded for rank 3 during the baseline or control time points (Suppl. Figure 3). In *A. perideraion*, there were no significant differences in cooperative behaviours between rank 4 in the treatment groups or rank 3 in the control groups at any time point (Suppl. Figure 3).

## Discussion

We show that there are differences in levels of aggression and cooperation among ranks in anemonefish size hierarchies, and that these behaviours are likely adaptive since they change in a predictable way as individuals ascend in rank. While species differed in overall levels of aggression recorded, we found consistently higher frequencies of aggression in higher ranks (more dominant individuals) in both species. Our results thus confirm predictions about increased aggression with higher rank and larger group size (Cant et al., 2006a) and are in line with findings in other fish organized in dominance hierarchies (Colleter & Brown, 2011; Dey et al 2013). However, our results do not confirm predictions that lower ranks should perform more cooperative behaviours (Cant & Field, 2001; 2005). Focussing just on the subordinates, we found the highest ranked subordinate (rank 3) performed most of the cooperative behaviours as well as aggressive behaviours. For cooperative effort, our results are very close to what is predicted when basal mortality rate is stable but the cost of helping effort increases for lower ranks (see Cant & Field, 2005). Since the dominance hierarchy in anemonefishes is organized by size, smaller ranks may not be able to bear the cost of cooperative behaviours, as has been found in some birds, mammals and other fishes (Boland et al., 1997; Clutton-Brock et al., 2000; Heinsohn & Legge, 1999). Taborsky (1984) demonstrated the cost of helping in a cooperatively breeding cichlid, by showing that subordinates performing cooperative behaviours grew slower than those that did not cooperate. In our study, smaller ranks spent more time performing submissive behaviours and the lowest ranks did not perform many of the behaviours observed in the higher ranks. Instead, they were seemingly occupied with feeding and avoiding other group members (T Rueger, pers. obs.), presumably in order to avoid eviction (Buston, 2003c) Thus, our results show the importance of studying aggressive and cooperative behaviours simultaneously and for each individual rank, and the importance of extending empirical testing to systems where individuals are not related.

We found some evidence for rank ascension within only a couple of days in both species of anemonefishes when the opportunity was experimentally created. Aggressive behaviours increased in rank 4 when rank 3 was removed and were comparable to those recorded for rank 3 in the control groups, where group size changes were accounted for. Also, while cooperative behaviours did not significantly increase, some upward trend was evident. The slight increase in cooperative behaviours, particularly cleaning and other interactions with the anemone, could be ‘prestige signalling’, similar to what has been recorded in the freshwater cichlid *Neolamprologus pulcher*, where the individual indicates its value by helping to improve territory quality (see review in Wong & Balshine, 2011). At the same time, the aggressive reaction of all group members towards the reintroduced fish (either rank 3 or rank 4) could be a punishment for leaving the group or being ‘idle’. Similar reactions to experimentally removed and reintroduced individuals have been recorded in *N. pulcher* (Balshine et al 1998; Fisher et al., 2014) and fairy wrens (Mulder & Langmore, 1993). Interestingly, aggression was directed from the rank 4 individuals towards the reintroduced, larger rank 3 individuals, suggesting that perceived rank ascension took place over a short time span (48hrs). However, since the experimental duration was insufficient to allow rank 4 individuals to grow and bridge the gap in total body length, initial protests appeared to be over quickly, with both rank 3 and 4 individuals present in their original ranks and aggressions subsiding 24hrs after reintroduction. Longer experiments involving growth rate measurements would be useful for understanding rank ascension and punishment structures in more detail. In any case, the current study shows that behaviours differ and change consistently among ranks and hence such variation is likely to be viewed as adaptive.

Our findings contribute to the growing body of literature disentangling the causes of sociality in groups of unrelated vertebrates. For anemonefishes, the reasons for subordinates to stay in groups of non-relatives and forego reproduction have been found to range from future fitness benefits through territory inheritance (Buston, 2004a), to the presence of social constraints (Buston, 2003c; Rueger et al., 2018) and environmental constraints (Branconi et al., 2020). However, why dominants tolerate unrelated, non-breeding subordinates has remained unclear, since subordinates have not been shown to directly increase dominant reproductive output, at least in the short term (Buston, 2004b). Here however, we provide evidence to show that subordinates do perform cooperative behaviours, and hence a reason why breeders tolerate them. These behaviours, namely the cleaning and massaging of the anemone and deterrent of anemone predators, are likely cooperative as they invoke a temporary decrease in subordinate fitness (time and energy expenditure that could be focused on other activities such as feeding) and yet provide indirect benefit to dominants. From the subordinate’s perspective, there is minimal relatedness among group members (Buston et al., 2007) and therefore no inclusive fitness to be gained through helping. Rather, cooperative effort is likely to be beneficial as it increases the quality of the territory (anemone) which in turn increases survival, access to resources and later inheritance of a higher quality territory (Kokko et al., 2001; Kingma et al., 2014). Since a substantial proportion of the cooperative behaviours observed in this study involved interacting with the anemone, it is likely that enhancement of territory quality is a key benefit of cooperation. In addition, since subordinates stand to inherit the territory upon death of the dominants, these cooperative behaviours directed towards the anemone could enhance the quality of the anemone territory, generating direct future benefits to helpful subordinates.

Interestingly, when comparing the two species that are closely related and share a very similar ecology, we found some distinct differences in the degree to which specific behaviours were displayed. The causes of higher aggression in *A. perideraion* compared to *A. percula* are not known but may be partially related to smaller size differences between the lower ranks *in A. perideraion* compared to *A. percula* (T Rueger, pers obs.). Size ratios are known to vary with respect to position in the hierarchy (Buston & Cant 2006; Wong, 2011). When lower ranks are closer in size, as is observed in *A. perideraion*, individuals may need to be more aggressive to protect their rank from those closer in size. While the cause of the interspecific differences in aggression observed here is unknown, future studies should consider the influence of size ratios on behavioural patterns in species with size-based dominance hierarchies. *A. perideraion* females have also been observed leaving large groups and forcefully taking over neighbouring anemones (Rueger et al., 2018), suggesting that elevated aggressiveness of *A. perideraion* could simply be a fixed trait of this species. Additionally, *A. perideraion* exhibited lower levels of cooperation (territory maintenance and defence) than *A. percula*, with more time being occupied with intragroup behavioural interactions. Higher levels of cooperation from subordinates in *A. percula* may indicate a more cohesive, efficient group structure and at the individual level could indicate that stricter conflict resolution mechanisms are at play. Future research should focus on the nuanced differences between species and explore the ecological factors that may lead closely related species of anemonefishes to pursue different social strategies and show different behavioural frequencies.

## Conclusion

Our interspecific comparative study has shown strong support for some theoretical predictions pertaining to variation in aggression levels between ranks in animal groups with dominance hierarchies, but we also found theory-contradicting results pertaining to levels of cooperative behaviours. This shows the importance of testing the generality of theory based on groups of relatives using animals living in groups of unrelated individuals. Another important aspect of this study was the quantification of subordinate behaviours that are helpful to the group and/or territory. Although we did not observe alloparental broodcare, we did find significant levels of cooperative behaviours in subordinates which are likely to have long-term impacts on the fitness of dominant breeders via effects on the anemone host. This contributes to the evidence expanding our view of sociality. Finally, we show that there are interspecific differences in aggressive and cooperative behaviours between closely related species. These differences warrant further investigation using broader comparative approaches in a wider range of species and likely represent the next frontier in sociality research.

## Acknowledgements

We thank the Tamare-Kilu communities for granting access to their reefs for this study. We thank Nelson Sikatua, Jerry Sikatua, Catheline Froehlich and Selma Klanten for field assistance and Chancey MacDonald for help with the artwork. Fieldwork was supported by Mahonia Na Dari Research and Conservation Centre, Kimbe Bay, Papua New Guinea.

## Ethics

This study was approved by the University of Wollongong Animal Ethics Committee, permit number AE18/06.

