## Supplemental Material for "Cooperative and aggressive behaviours vary between ranks in anemonefish social hierarchies"

Preprint

**Ethogram**

**Suppl. Table 1.** Ethogram adapted for the purpose of this study from Wong et al. 2013.

| Behavioural category | Behaviour | Description |
|---|---|---|
| Aggressive | Chases | Fish rapid swims towards conspecific and conspecific flees. |
| | Bites | Fish bites another fish. |
| | Jolts | Rapid darts and turns prior to or during aggressive contact. |
| | Aggressive display | Fish meets another, turns side on with erect fins, stiffened body posture. |
| Submissive | Flee | Fish rapidly swims away after being chased or bitten. |
| | Body shake | Fish shakes body side to side in jolty fashion. |
| Cooperative | Bite anemone | Fish bites mouth or tentacles of anemone. |
| | Massage anemone | Fish moves rigorously against anemone foot (often observed when anemone is contracting). |
| | Cleaning | Fish picks up sand or debris and spits it out outside anemone. |
| | Defense against predator | Quick swims or bites directed towards heterospecifics, either potential egg predators (e.g., wrasses), or anemone predators (e.g. butterflyfishes). |
| | Defense against competitor | Quick swims or bites directed towards heterospecific food competitors (e.g., damselfishes) |
| | Parental | Fish is within one body length of the nest. May be fanning or mouthing eggs. |

| | | |
|---|---|---|
| Neutral | Meeting | Fish simultaneously approach each other to within 1 body length (may be followed by soft touch). |
| | Soft touch | Fish make contact with fins or body before separating again, preceded by meeting. |
| | Follows | Fish follows another fish within 1 body length for at least a few seconds. |
| Out of sight | | Fish cannot be seen because it enters anemone, moves under or behind anemone, or swims beyond the video frame. Measured in seconds. |

Preprint

**Rank ascension experiment results- Post-hocs and contrasts**

**Aggression**

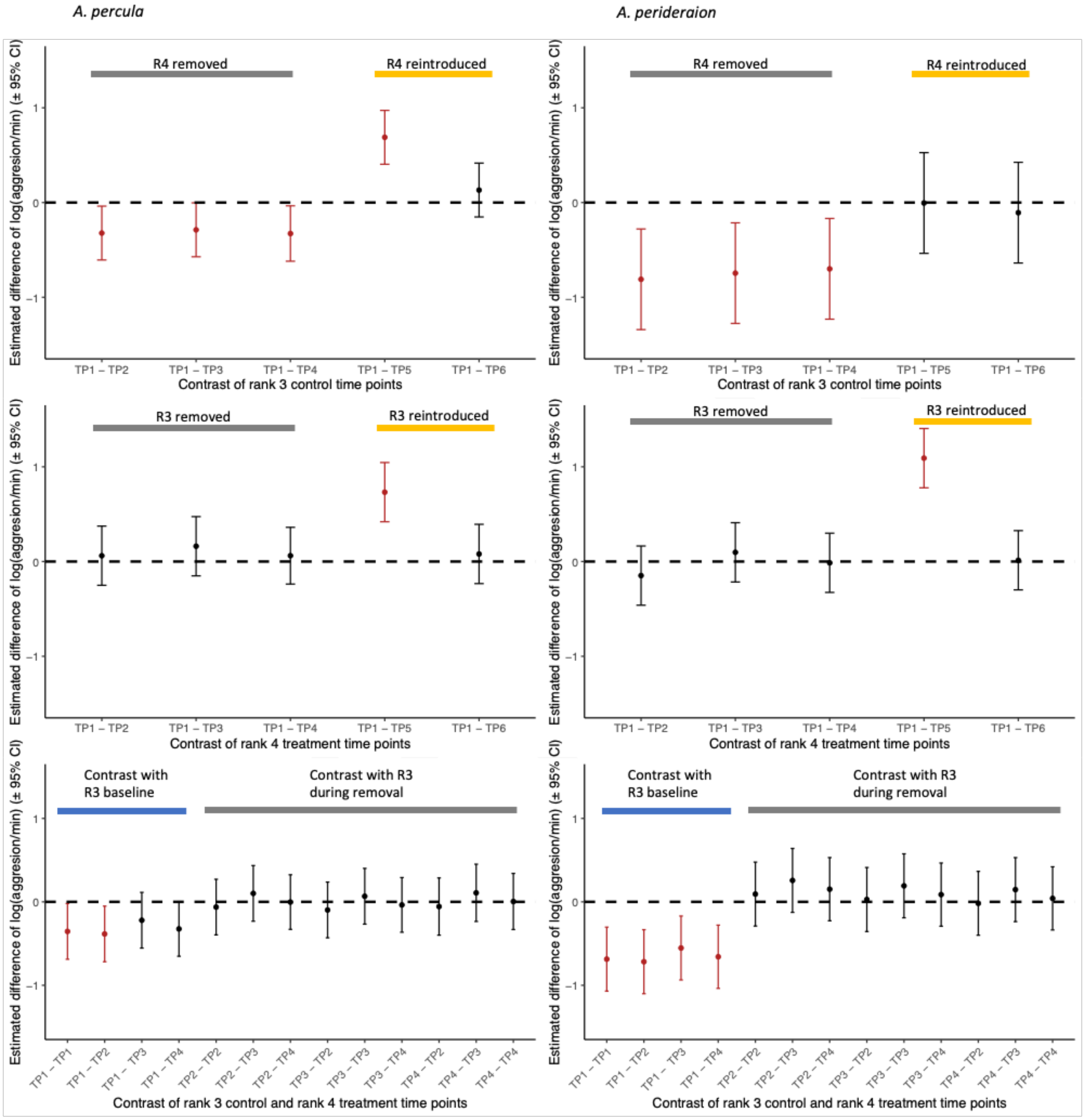


**Suppl. Figure 1.** Post-hoc comparisons and contrast for aggressive behaviours in *A. percula* and *A. perideraion*. Row one: Estimates (+/- 95% confidence intervals (CI)) of differences in rank 3 aggressive behaviours (log aggression/min) between the baseline time point (TP1) and each time point during the control (TP 2-6). Row two: Estimates (+/- 95% confidence intervals (CI)) of differences in rank 4 aggressive behaviours (log aggression/min) between the baseline time point (TP1) and each time point during the treatment (TP 2-6). Row 3: Estimates (+/- 95% confidence intervals (CI)) of differences in aggressive behaviours (log aggression/min) between rank 3 control time points (baseline TP1, rank 4 removed TP2-4) and rank 4 treatment time points (baseline TP1, rank 3 removed TP2-4). Red indicates contrast is statistically significant ($p < 0.05$).

### Submission

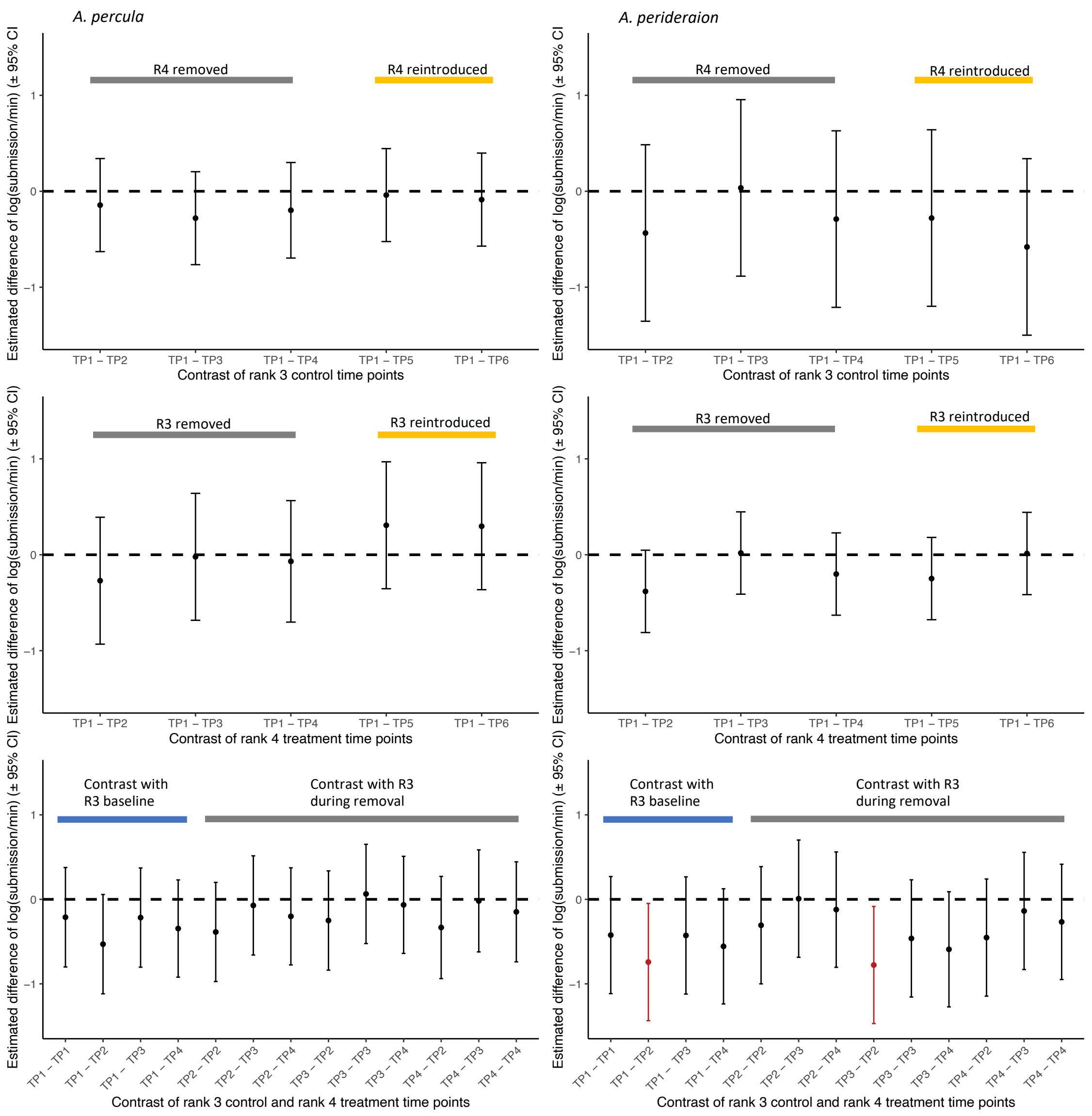


**Suppl. Figure 2.** Post-hoc comparisons and contrast for submissive behaviours in *A. percula* and *A. perideraion*. Row one: Estimates (+/- 95% confidence intervals (CI)) of differences in rank 3 submissive behaviours (log submission/min) between the baseline time point (TP1) and each time point during the control (TP 2-6). Row two: Estimates (+/- 95% confidence intervals (CI)) of differences in rank 4 submissive behaviours (log submission/min) between the baseline time point (TP1) and each time point during the treatment (TP 2-6). Row 3: Estimates (+/- 95% confidence intervals (CI)) of differences in submissive behaviours (log submission/min) between rank 3 control time points (baseline TP1, rank 4 removed TP2-4) and rank 4 treatment time points (baseline TP1, rank 3 removed TP2-4). Red indicates contrast is statistically significant ($p < 0.05$).

### Cooperation

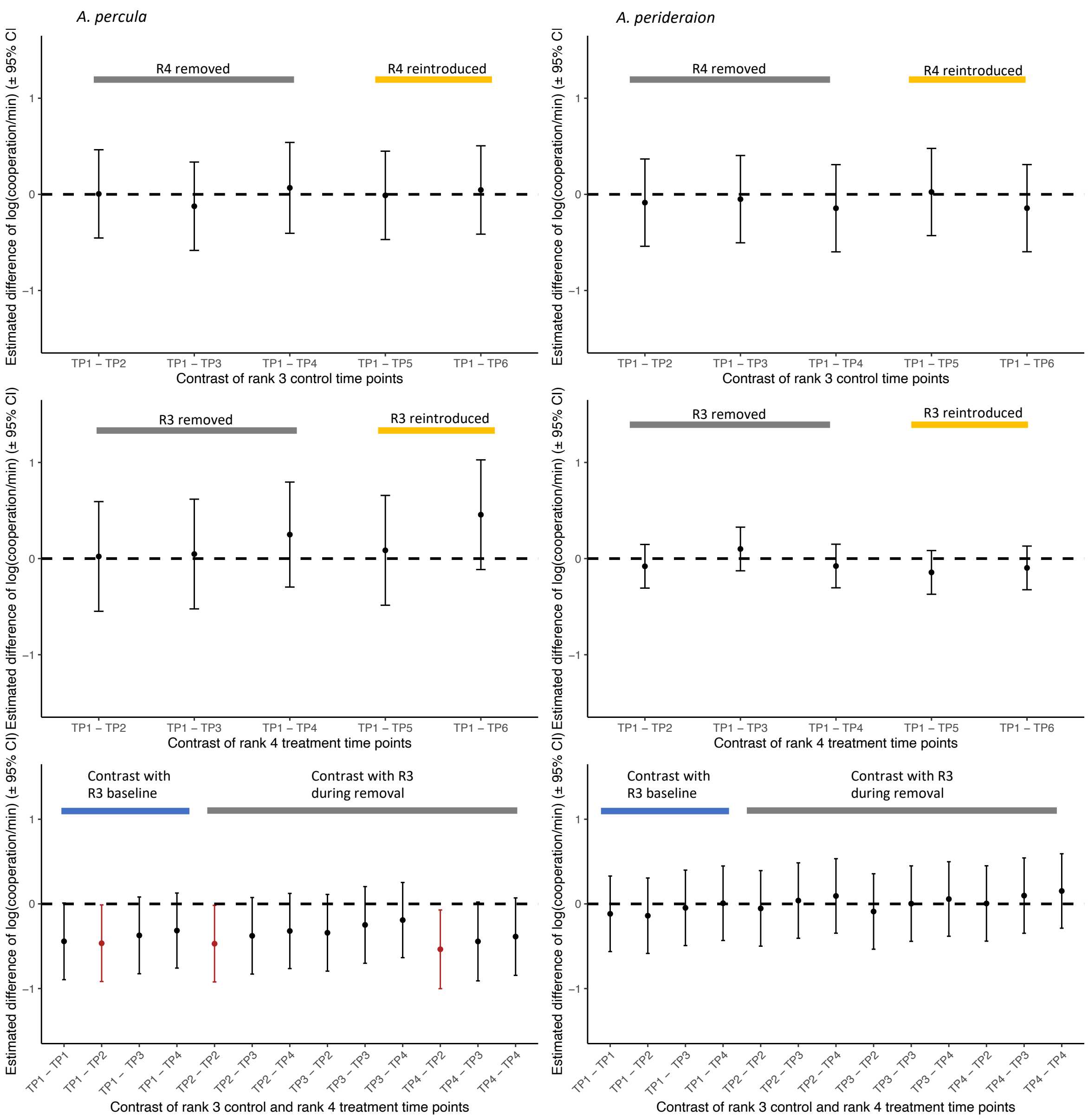


**Suppl. Figure 3.** Post-hoc comparisons and contrast for cooperative behaviours in *A. percula* and *A. perideraion*. Row one: Estimates (+/- 95% confidence intervals (CI)) of differences in rank 3 cooperative behaviours (log cooperation/min) between the baseline time point (TP1) and each time point during the control (TP 2-6) . Row two: Estimates (+/- 95% confidence intervals (CI)) of differences in rank 4 cooperative behaviours (log cooperation/min) between the baseline time point (TP1) and each time point during the treatment (TP 2-6). Row 3: Estimates (+/- 95% confidence intervals (CI)) of differences in cooperative behaviours (log cooperation/min) between rank 3 control time points (baseline TP1, rank 4 removed TP2-4) and rank 4 treatment time points (baseline TP1, rank 3 removed TP2-4). Red indicates contrast is statistically significant ($p < 0.05$).

Preprint
